# Single-cell resolution unravels spatial alterations in metabolism, transcriptome and epigenome of ageing liver

**DOI:** 10.1101/2021.12.14.472593

**Authors:** Chrysa Nikopoulou, Niklas Kleinenkuhnen, Swati Parekh, Tonantzi Sandoval, Farina Schneider, Patrick Giavalisco, Mihaela Bozukova, Anna Juliane Vesting, Janine Altmüller, Thomas Wunderlich, Vangelis Kondylis, Achim Tresch, Peter Tessarz

## Abstract

Epigenetic ageing clocks have revealed that tissues within an organism can age with different velocity. However, it has not been explored whether cells of one type experience different ageing trajectories within a tissue depending on their location. Here, we employed lipidomics, spatial transcriptomics and single-cell ATAC-seq in conjunction with available single-cell RNA-seq data to address how cells in the murine liver are affected by age-related changes of the microenvironment. Integration of the datasets revealed zonation-specific and age-related changes in metabolic states, the epigenome and transcriptome. Particularly periportal hepatocytes were characterized by decreased mitochondrial function and strong alterations in the epigenetic landscape, while pericentral hepatocytes – despite accumulation of large lipid droplets – did not show apparent functional differences. In general, chromatin alterations did not correlate well with transcriptional changes, hinting at post-transcriptional processes that shape gene expression during ageing. Together, we provide evidence that changing microenvironments within a tissue exert strong influences on their resident cells that can shape epigenetic, metabolic and phenotypic outputs.

## INTRODUCTION

Ageing is characterised by a general physiological decline that is accompanied by metabolic, epigenetic and transcriptional changes^1^. A common attribute for these alterations is an increased inter-individual heterogeneity as observed in large cohorts. Even on an organismal level within populations of genetically identical individuals, variability seems intrinsically inter-connected with ageing. For example, in cohorts of *C. elegans* or mice, some individuals die much earlier than others^2^.

It is largely appreciated that transcriptional variability increases with age^3–5^. While whole tissue omics approaches have been important to get an insight into the uniform changes that occur on the organ level during ageing, such methods cannot investigate heterogeneity on a cellular level. It is therefore unresolved whether all cells of the same cell type in a tissue age in the same way or whether the location of the cells within a tissue matters in this context. The development of single-cell and spatial omics methods renders it now possible to obtain (spatially resolved) molecular profiles at close to single-cell resolution, thus providing promising tools for deciphering the multifaceted process of ageing^6^.

The liver is a heterogeneous tissue that consists of hepatocytes arranged in repeating units of hexagonally shaped lobules. Blood flows into the lobule from portal veins and hepatic arteries at the corners of the lobules to the central veins. This architecture creates gradients of oxygen, nutrients and hormones^7^. This gradual change in the lobule’s microenvironment is also referred to as liver zonation^8^ and the resulting spatial division of labour is essential for the optimal function of the liver. For example, the outer highly oxygenated periportal lobule layers perform mitochondrial-dependent metabolic tasks such as *β*-oxidation whereas the low oxygen concentrations at the pericentral areas will drive glycolysis^7^. As hepatocytes are the primary cells that perform these metabolic processes and their metabolic characteristics depend on location, the liver is an attractive tissue to address the impact of location and metabolic state on the ageing trajectory within a dedicated cell type.

Here, we employed spatial transcriptomics as well as single-cell ATAC-seq (scATAC-seq) in conjunction with publicly available single-cell RNA-seq (scRNA-seq) data from ageing mice to address how ageing of hepatocytes is affected by zonation in the liver. One very obvious phenotypic difference in the ageing and diseased liver is the deposition of fat, which is mainly focused around the central vein. Using spatial transcriptomics, we report insights into the molecular underpinnings of this phenotype, and additionally identify mitochondrial dysfunction as a potential driver for age-related phenotypes in the periportal region of the liver. While scATAC-seq can clearly separate young and old hepatocytes, unsupervised clustering approaches do not separate scRNA-seq profiles based on their age. Yet, age is a relevant factor for explaining transcriptional variability between cells. Together, the data presented here shed light on the molecular basis of fat deposition in the ageing liver and serve as a valuable resource for the hepatic and ageing community.

## RESULTS

### Spatial Transcriptomics give insights into the zonation-specific and age-related metabolic rearrangements

Transcriptional profiling using bulk RNA sequencing data from the Tabula Muris Consortium^9^ shows metabolic pathways, known to be changing in ageing^10^, with the majority of genes contributing to alterations in lipid metabolism (Figure S1a,b, Supplementary Table 1). Changes in lipid metabolism have been described to occur during ageing and the recent development of lipidomics started to identify corresponding changes in lipid profiles^11^. Liver pathologies that involve fat deposition, such as non-alcoholic fatty liver disease (NAFLD) show a tendency towards zonated lipid deposition around the central area^12^, but we were not aware of any dataset investigating lipid deposition in the ageing liver with respect to the specific zones. To assess the lipid deposition around the main zones, we performed RNAScope for pericentral (Cyp2e1, Glul) and periportal markers (Albumin, Cyp2f2)^13^ combined with H&E (Hematoxylin and Eosin) staining in liver isolated from young (3-4 months) and old (18-20 months) mice (Figure 1a, S1c). Importantly, Sirius red staining showed no profound increase in liver fibrosis in old livers (Figure S1c). On the contrary, Oil-red-O (O-R-O) staining (Figure S1d, upper panel) and immunohistochemical (IHC) staining for PLIN2 (Figure S1d, lower panel), a protein known to be enriched at the outer membrane of LDs^14^, showed that large LDs accumulate around the central vein in aged livers.

**Figure 1:**
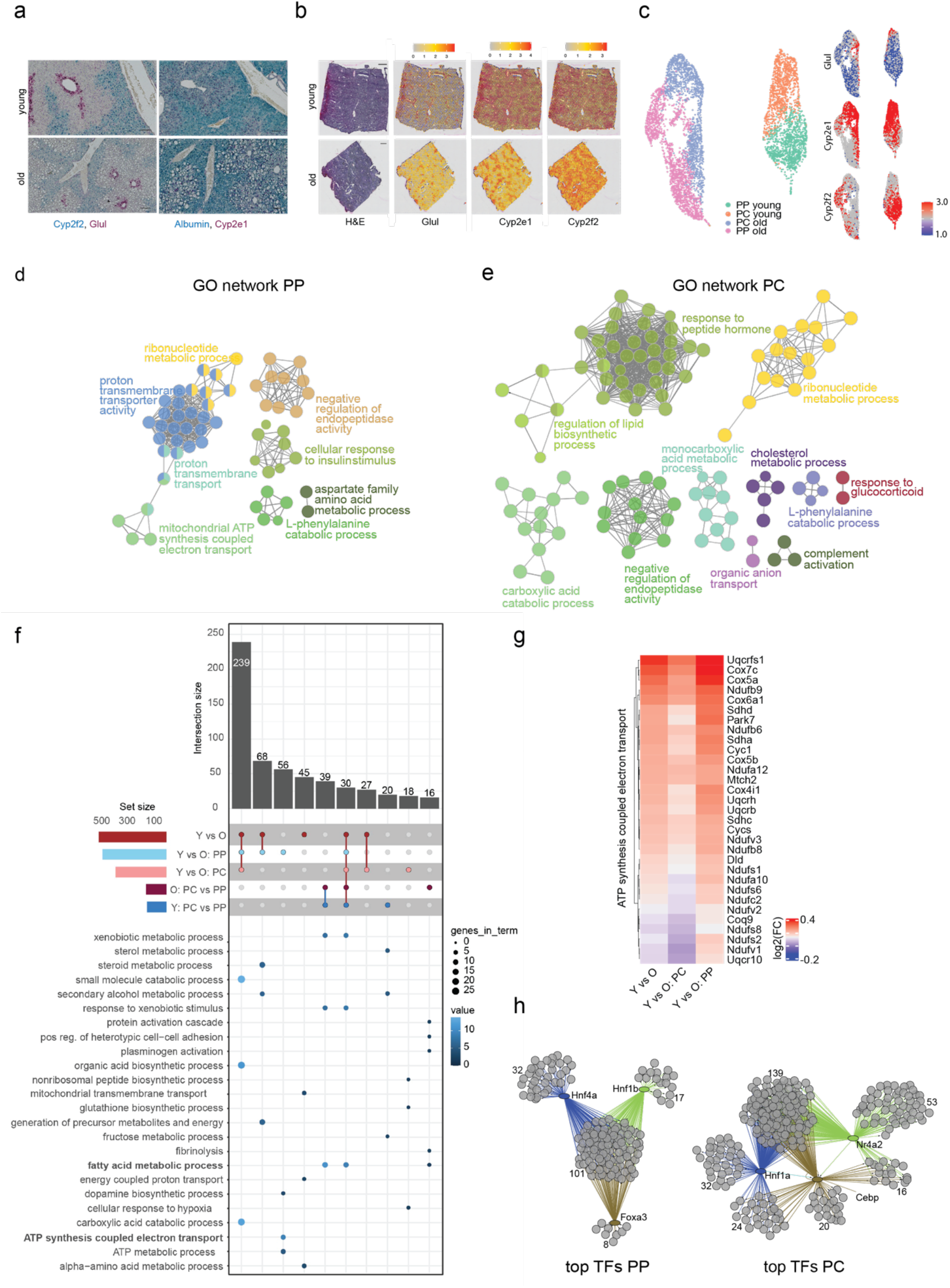
Age-related and zonation-specific transcriptional alterations. a) RNAscope of zone-specific marker genes Glul (magenta, upper panel), Cyp2f2 (cyan, upper panel), Cyp2e1 (magenta, lower panel) and Albumin (cyan, lower panel) in paraffin-embedded liver sections from young (3 month old) and old (18 month old) mice. Scale bar = 100 µm. b) H&E staining of one young (upper panel) and one old (lower panel) liver specimen used for spatial transcriptomics (Scale bar=500 µm) and plots showing the expression levels of Glul, Cyp2f2 and Cyp2e1 indicated by colour. The colour gradient represents normalised gene expression. c) UMAP projection of the spatial data, colour-coded are the different zones and ages (left panel) and the expression of Glul, Cyp2e1 and Cyp2f2 (right panel). d), e) GO network calculated using ClueGO for differentially expressed genes in the periportal (based on Supplementary Table 3 - for details, see Method section) (d) and pericentral (e) zone of the ageing liver. f) UpSet plot showing the number of differentially regulated genes (top) and pathways (bottom) in the indicated categories (Y=young, O=old, PC=pericentral, PP=periportal). g) Heatmap with hierarchical clustering of differentially expressed genes from the indicated pathways selected from f). g) Transcription factor activity prediction from the age-dependent differentially expressed genes by the iRegulon app in Cytoscape (based on Supplementary Table 3 - for details see Methods section). For each zone, the top predicted TFs are shown as well as their interaction to regulate transcripts. Numbers indicate the genes in every cluster.

The apparent zone-dependent deposition of lipids in the ageing liver prompted us to investigate the underlying transcriptional events. We used the 10X Genomics Visium Platform and ran 10µm tissue cryosections from livers of two young and two old mice. The sequencing metrics of the samples can be found in Supplementary Table 2. Initially, we visualized the normalized spatial gene expression of the zonation markers Cyp2f2, Cyp2e1 and Glul in young and old liver (Figure 1b). Based on the expression distribution of these marker genes, spatial transcriptomics was able to resolve central and portal areas.

Principal Components analysis showed that spots from each slide cluster; spots from the two young liver slides overlap, while spots from the two old slides separate (Figure S1e). Therefore, to guard against batch effects, we integrated young and old datasets individually, first using canonical correlation analysis^15^ and analysing zonal expression effects. Then, we merged all datasets using the same strategy. To assess whether the sample separation reflected gene expression differences based on age or were mostly due to a potential batch effect, we used the loadings calculated in the PCA and intersected those with a recently published resource, in which global ageing genes were defined organismal and tissue-wide^16^. The majority of the genes that contributed to the first principal component were part of the liver-specific global ageing genes (Figure S1f). To perform differential analysis of the PP and PC zones of the liver tissue, we assigned spots to pericentral and periportal groups based on Cyp2e1 and Cyp2f2 expression levels (Figure 1c, Methods). We used a two-part, generalised linear hurdle model^17^ to identify gene expression changes between young and old liver in general, but also specifically in periportal and pericentral region upon ageing (Supplementary Table 3). We performed GO enrichment using the ClueGo plugin for Cytoscape^18, 19^ for the age-related changes in the two zones. While the periportal region was characterized by changes in mitochondrial respiration and proton transport as well as amino acid metabolism (Figure 1d), ontologies in the pericentral zone were enriched for terms related to lipid biosynthesis and carboxylic acid catabolic processes (Figure 1e). Common for both zones were changes in ribonucleotide metabolism and response to peptide hormones, such as insulin (Figure 1d,e). To further zoom into the differences of the zones, and to identify commonly and zone-specifically deregulated genes, we represented the data as an UpSet plot (Figure 1f). This analysis confirmed the notion of zone-specific alterations. The periportal area showed age-related expression changes of genes encoding for members of the electron transport chain, for example an age related decrease in Uqcrfs1 (cytochrome b-c1), which catalyses the electron transfer from ubiquinol to cytochrome c^20^, and Cox7c or Cox5a that drive oxidative phosphorylation^21^ (Figure 1g). On the other hand, the pericentral area showed a signal of hypoxia, which might be caused by the previously reported changes in liver vascularisation upon ageing^22^. Finally, we wanted to understand whether the transcriptional changes were driven by a dedicated set of transcription factors. We used the iRegulon app within Cytoscape^18, 23^ and visualised the top three most significant TFs (NES >4) based on age-dependent differential expression within the two zones. Shared between the zones is Hnf1, which has been shown to regulate many hepatic genes^24^. Genes in the periportal area were predicted to be regulated by Hnf4a and Foxa3 (Figure 1h). Hnf4a is a master regulator during hepatic differentiation and plays an important role during liver regeneration^25^, similarly to Foxa3^26^. In addition, Hnf4a has recently been shown to possess anti-proliferative capacity and thus protects against hepatocellular carcinoma^25^. On the other hand, genes in the pericentral zone were predicted to be regulated by Cebp and Nr4a2 (Figure 1h), two TFs that regulate glucose and lipid metabolism^27, 28^. Taken together, spatial transcriptomics revealed that ageing is accompanied by zonation-specific metabolic rewiring, which is driven by a network of dedicated transcription factors.

### The ageing liver is characterised by lipid remodelling and loss of spare respiratory capacity in periportal mitochondria

The spatial transcriptomic data suggested age-related metabolic alterations that depend on the location of cells with respect to central or portal regions. To gain more insight into the metabolic alterations, we first performed lipidomics to characterise the changes in lipid metabolism within the ageing liver. This approach allowed us to address not only storage and membrane lipids, but also to analyse levels of cardiolipins and ubiquinones to further investigate the observed alterations in mitochondrial metabolism.

We extracted lipids from livers of young and old mice. PCA (Figure S2a) and differential abundance analysis of the most significantly changed lipids (Figure 2a, Supplementary Table 4) showed a strong lipid remodelling for most of the major lipid classes. While we did not observe an overall increase in triacylglycerides (TAGs), we noted a significant increase in the levels of lysoPE (LPE) and lysoPC (LPC) (Figure 2b), which might stem from the remaining serum in the liver as those lipids are enriched in extracellular fluids ^30^. Importantly, we noted a strong increase in diacylglycerides (DAGs) and a decrease in sphingomyelin (SM) (Figure 2b), pointing towards changes in membrane fluidity^31, 32^ and lipid-mediated signalling. Indeed, an increase in DAGs as well as a decrease in SMs has been linked to an increase in insulin insensitivity, a well-known hallmark of ageing^33, 34^ and a pathway that was also evident in the spatial transcriptomics data (Figure 1d,e). We then focused on mitochondria-related lipids. A significant increase in all cardiolipins (CL) measured (Figures 2b, S2b) indicated changes in the composition of mitochondrial membranes and hence the function of mitochondrial inner membrane proteins, including the electron transport chain (ETC)^35^. This hypothesis was also supported by the observation that ubiquinones, lipids that transfer the electron between the different complexes of the ETC, were strongly down-regulated with age (Figure 2c). These findings in combination with the spatial transcriptomics data supported the hypothesis of age-dependent mitochondrial changes. As the spatial transcriptomic data and the lipidome analysis pointed towards a strong impact on mitochondrial metabolism, we wanted to investigate this phenotype in more detail, particularly in a zone-specific manner. In order to do this, we used a previously published protocol^36^ to sort hepatocytes into pericentral and periportal upon perfusion of the liver (Figure 2d, S2c). This approach depends on the zonation-dependent expression of E-cadherin (periportal) and CD73 (Nt5e, pericentral)^36^ and was able to separate pericentral and periportal hepatocytes as judged by expression of Glul and Cyp2f2 (Figure S2d). First, we measured mitochondrial content in the two zones in an age-dependent manner, which was variable across different animals and zones, but largely unaltered with age (Figure S2e). Finally, we performed Seahorse analysis using the mitochondrial stress kit to assess mitochondrial function. While basal respiration and ATP production changed only mildly with age, we observed a striking reduction in the maximal and thus, spare respiratory capacity (SRC) in periportal hepatocytes (Figure 2e). On the other hand, pericentral hepatocytes showed an increase in maximal respiration. Loss of SRC sensitizes the cells to surges in ATP demand^37^ and it has been proposed that SRC can be used as a measure of mitochondrial health^38^. Taken together, spatial data, lipidomics and bioenergetics measurements point towards an age-dependent decrease in hepatic mitochondrial fitness and function, specifically in the periportal zone of the liver.

**Figure 2:**
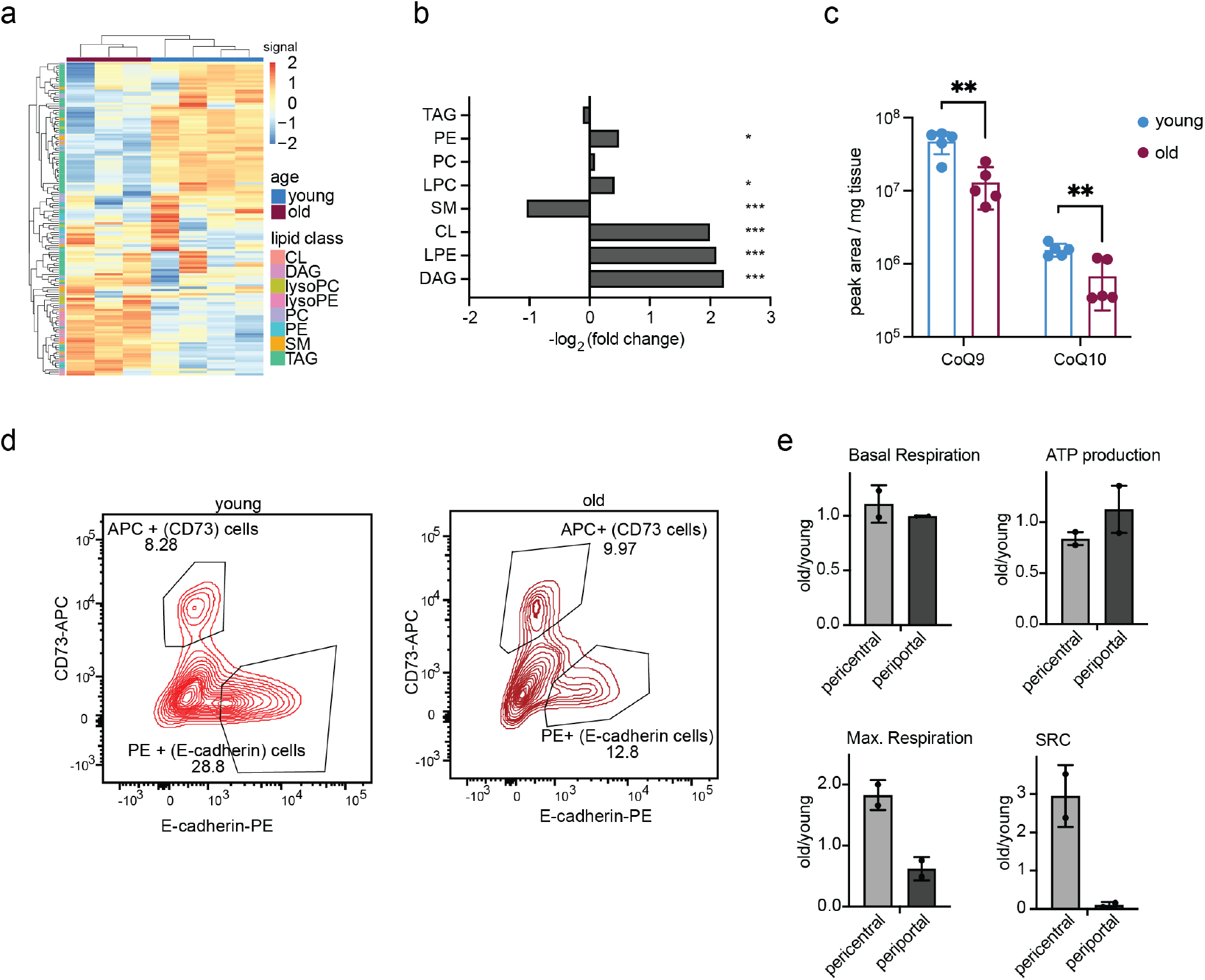
Lipid remodelling and alterations in mitochondrial metabolism in the ageing liver. a) Heatmap with hierarchical clustering of lipid datasets derived from 3 old and 4 young mouse livers, showing the differentially expressed classes of lipids. Hierarchical clustering was performed using LipidSig ^29^ based on data available in Supplementary Table 4. b) Bar plot of the log-fold changes in lipid classes expressed in old vs. young liver. Fold changes and significance (*p-value<0.05, ***p-value<0.001) were calculated using LipidSig based on data available in Supplementary Table 4. c) Bar plot showing the expression of Ubiquinones CoQ9 and CoQ10 in young and old liver. Statistical significance was determined using a two-sample t-test (**p-value<0.01). d) Exemplary FACS profiles of sorted hepatocytes based on CD73 (pericentral) and E-Cadherin (periportal). e) Mitochondrial function as measured by Seahorse Mitochondrial Stress kit (parameter on top of graph) expressed as old vs young and pericentral-periportal. N=2 (per N, one or two young and two old mice were sorted and averaged). Error bars represent the SEM.

### Chromatin accessibility in mouse liver carries a hepatocyte ageing signature

Having defined the transcriptional, lipid and functional alterations that occur within the periportal and pericentral zones of the ageing liver, we next wanted to investigate if the differences in phenotype and transcriptome might be explained by an underlying change on the epigenetic level. Therefore, we performed scATAC-seq using the 10x Chromium platform. We profiled 4838 nuclei prepared from three young liver tissues and 3361 nuclei from three old liver tissues. Sequencing metrics can be found in Supplementary Table 3. In order to identify cell types and their accessibility profiles, we combined the young and old datasets and subsequently analysed them together using cisTopic^39^. Clustering according to cell-to-cell similarity using UMAP identified several cell clusters. Most of the clusters showed intermixing between young and old cells. However, the biggest cluster showed a clear separation between the two age groups (Figure 3a).

**Figure 3:**
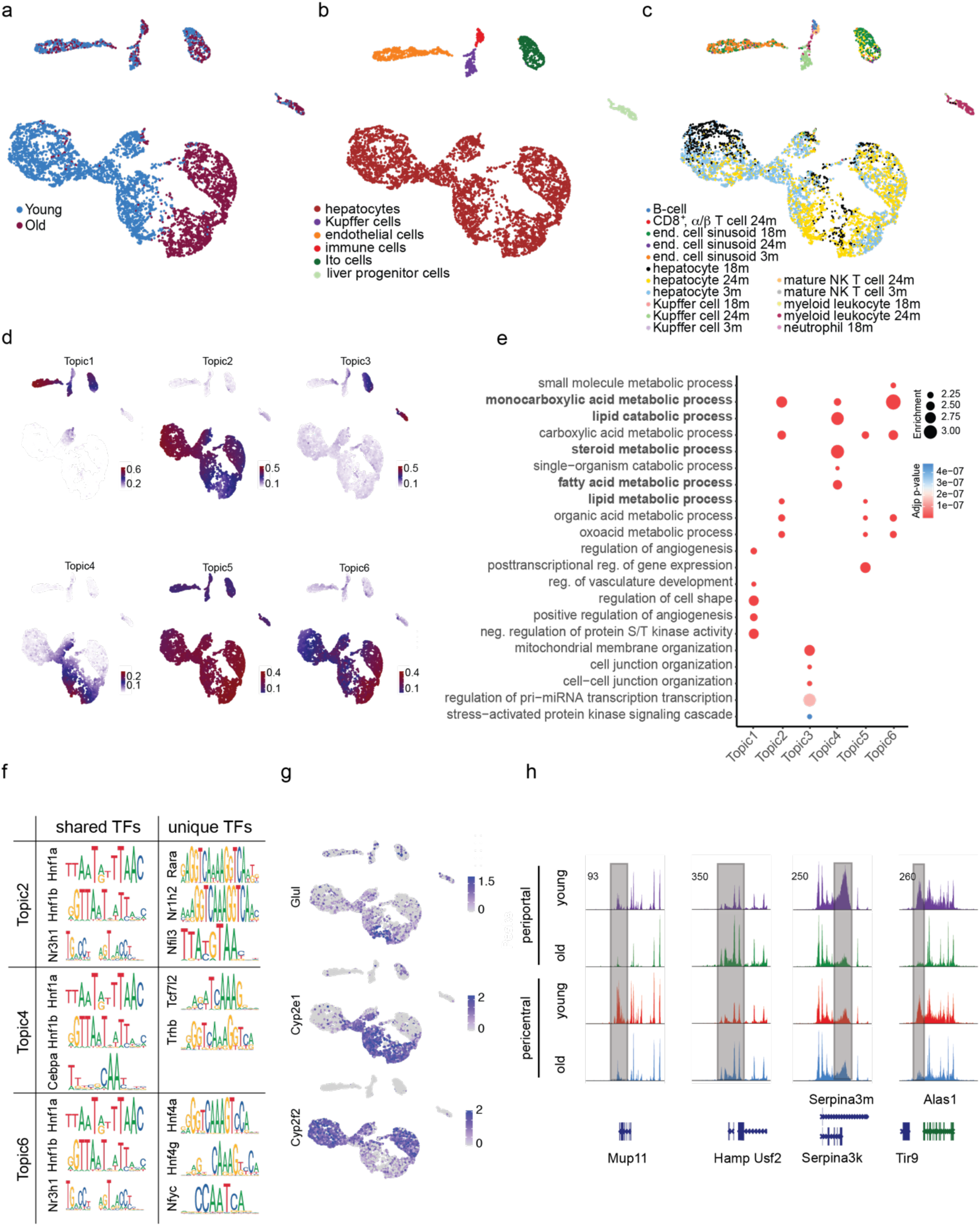
Differential chromatin accessibility in aged liver hepatocytes. a-c) UMAP projection of scATAC-seq data of mouse liver nuclei. a) Different colours represent liver cells from young and old age groups identified using cisTopic. b) Different colours represent different cell types based on imputed marker gene activity (see also Supplementary Figure S3C). c) Different colours represent different cell types predicted with cell type assignment using the FACS data of the TMS ^9^ d) cisTopic identified six different topics. Colour code of the UMAPs is according to the normalised topic score for each cell. e) GO term analysis of the 6 different topics. Highlighted are liver-associated metabolic processes. f) Shared and unique transcription factor (TF) motifs corresponding to the ‘’hepatocyte’’ topics 2, 4 and 6. g) UMAP projections as in A. Colour code corresponds to the imputed gene activity of zone-specific genes Glul, Cyp2e1 (pericentral) and Cyp2f2 (periporal). h) Exemplary tracks of differentially accessible sites between pericentral and periportal hepatocytes upon ageing. The grey bar indicates altered regions.

This behaviour was confirmed by a complementary clustering using Signac^40^ (Figure S3a). To identify cell types, we inferred transcriptional activity from the respective promoter accessibility, as described previously^41^. We used known marker genes^13, 42^ and CellMarker (http://bio-bigdata.hrbmu.edu.cn/CellMarker/) to infer the cellular identity of each cluster. We were able to resolve all expected cell types of the liver, except for cholangiocytes (Figure 3b, S3b-c). We were not able to distinguish different immune cell types since their marker genes’ imputed activity was ambiguous (Figure 3b, S3b,c). In line with the observation that the livers were not fibrotic, we did not observe a significant increase in immune or hepatic stellate cell numbers based on the scATAC-seq profiles or detected a specific inflammatory signal. Notably, based on the marker gene profiles, the only cluster clearly separated by age was the hepatocyte one (Figure 3a,b, S3a,b). Regions that changed accessibility with age encoded for genes involved in pathways such as glucose homeostasis and fat-cell differentiation (Figure S3d). To further validate our chromatin-state-based cell type assignment, we predicted cell types of our scATAC-seq data with FACS-based scRNA-seq (Smart-seq2) data from the TMS consortium^9^. The integration largely confirmed our cell type prediction (Figure 3c). However, we noticed that particularly hepatocytes were not predicted clearly in different age groups.

Next, we made use of the inferred *cis*-regulatory topics that underlie the Latent Dirichlet Allocation (LDA) used by cisTopic^39^ and assigned those topics to the individual clusters. Most topics referred to specific cell clusters. Four topics were enriched in the hepatocyte cluster (Figure 3d). In line with the predicted cell types, GO terms associated with the hepatic topics centred around lipid and xenobiotic metabolism, the epithelial topic around angiogenesis and vasculature development, whereas the other topics were mostly associated with regulatory terms (Figure 3d). Interestingly, topics 2 and 6 correspond to young and old hepatocytes, respectively, whereas topic 4 was shared between the two age groups. Topics were further exploited to predict enriched transcription factor motifs. Here, we particularly focused on the three hepatic topics (Figure 3f, Supplementary Table 5). In topics 2, 4 and 6 well-known hepatic transcription factors were predicted, such as Hnf1a,b (see also Figure 1h). Each topic also contained its unique set of transcription factors that were specifically predicted to topic-defining regions. In topic 2, which was enriched predominantly in the young hepatocytes, we identified unique TFs to be Nr1h2, which is involved in steroid metabolism as well as Nfil3, which controls Per1 and Per2 and is thus involved in circadian rhythm. Recent work has highlighted the importance of the circadian clock during the ageing process, and changes in the clock dynamics are particularly altered in the ageing liver^43^. The shared topic 4 was characterised by TFs involved in b-catenin and Wnt signalling, Tcf7l2 and Trhb, which is linked to b-catenin production through thyroid signalling^44^. Finally, topic 6, which is enriched in old hepatocytes, contained Hnf4a as a predicted unique TF. These unique transcription factors predicted for each of the topics implied very specific regulation of metabolic and signalling pathways with age. In general, the enriched transcription factor motifs were in good agreement with the prediction based on the zone-specific and age-dependent differential expression (Figures 1h and 3f, Supplementary Table 5). The apparent age-dependent separation between topics 2 and 6 and their respective enrichment in young or old liver prompted us to investigate whether liver zonation might be associated with the topics’ separation. To test this, we imputed the gene activity of Glul, Cyp2e1 and Cyp2f2. Remarkably, there is a very clear separation in the scATAC-seq feature plots (Figure 3g). Using the apparent activity level of these three marker genes, we concluded that topic 4 represented the pericentral region, whereas topic 2 described the chromatin state for young periportal hepatocytes and topic 6 encompassed mostly old hepatocytes. The loss of a clearly defined periportal cluster is interesting and might be connected to the change in mitochondrial metabolism. Changes in mitochondrial metabolism have been shown to perturb stem and somatic cell function in ageing^45, 46^ and may lead to dysfunction of hepatocytes and other resident liver cells in the periportal area. The differences in accessibility between the zones with respect to peak enrichment can also be seen in other representative gene loci (Figure 3h). Taken together, scATAC-seq is able to reveal changes in the epigenome of single-cells and can resolve zonation-specific differences in chromatin states.

### Specific Cidea expression in the periportal zone is driven by chromatin architectural changes

How do chromatin alterations connect to the transcriptional program to drive age-related phenotypes? To address this question in more detail, we initially inspected the differentially expressed genes (periportal - 544; pericentral 429) that were changed with age. Intriguingly, we identified two members of the Cide gene family (Cidea and Cidec, or Fsp27) to be upregulated specifically in old pericentral hepatocytes (Figure 4a,b). Cideb on the other hand was expressed across both ages and zones. We used this gene family as paradigm to understand the connection between chromatin, transcription and phenotype as the expression showed a very clear distribution. In addition, all three Cide proteins have been shown to bind to LDs and to modulate LD dynamics^47–49^. Overexpression of Cidec in hepatocytes was sufficient to generate large LDs^48, 50^ and using electron microscopy, we found that the median size of LDs increased 4-fold with age (Figure 4c), which correlated well with the increased pericentral expression of Cidea and Cidec. We then turned to our scATAC-seq dataset and probed whether there was an underlying alteration in accessibility at the Cidea locus, potentially explaining the increase in expression. Indeed, we observed a specific age-dependent increase in accessibility at the Cidea locus (Figure 4d). Co-accessibility analysis using Cicero^51^ also identified the enhanced usage of a potential intronic enhancer within Cidea as marked by H3K27ac (Figure 4d).

**Figure 4:**
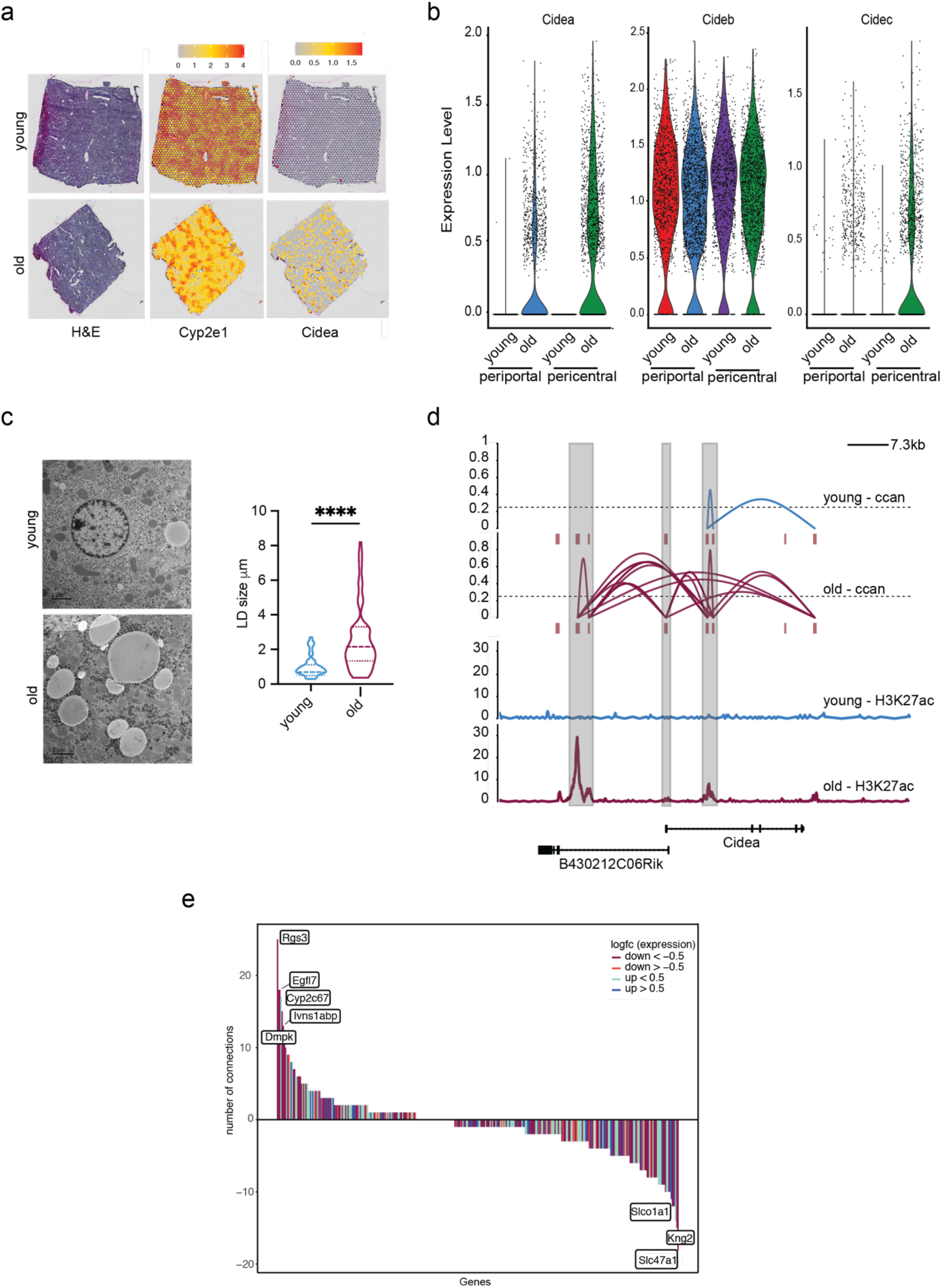
Connection between chromatin and transcriptional alterations in the ageing liver. a) H&E staining of one young (upper panel) and one old (lower panel) liver specimen used for spatial transcriptomics and a plot showing the expression level of Cidea. Please note that H&E stain and Cyp2e1 plots are identical to Figure 1b and used here for reference only. The colour gradient represents normalised gene expression. b) Violin plots indicating the expression levels of Cidea, Cideb and Cidec across pericentral and periportal regions in young and old liver. c) Transmission electron micrograph of lipid droplets (LDs) of young and old liver tissue. Representative images at 3000x, scale bar = 2 µm. ImageJ quantification of the mean LD diameter size in μM from ten randomly selected photos from a young (LD n=104, mean=0.8771) and ten from an old (LD n=88, mean=2.611) mouse specimen. Statistical significance was determined using an unpaired two-tailed t-test; ****p-value<0.0001. d) Ccan values based on Cicero ^51^ prediction of co-accessibility (upper panel) and the enhancer mark H3K27ac (lower panel) at the Cidea locus in young and old mouse liver. Highlighted in grey are potential enhancer and promoter regions from Cidea and its associated antisense long non-coding RNA, respectively. e) Age-related changes in co-accessibility of loci identified using spatial transcriptomics. Y-axis shows the differences in predicted contact points between young and old hepatocytes. Colour of the graphs highlight direction of gene expression change as taken from the spatial transcriptomics data (Supplementary Table 3) between young and old.

Given the apparent correlation between locus opening, potential enhancer engagement and transcription output at the Cidea locus, we next asked whether changes in co-accessibility might be a good predictor for differential gene expression on a global scale. We used the list of 482 differentially expressed genes between young and old and calculated the difference in chromatin accessibility for those genes (Figure 4f, Figure S4). In line with previous reports ^52^, we did not detect a general correlation between an increase in co-accessibility and transcription, indicating that co-accessibility is not a determinant for transcription. We noted as well that in many cases the levels of H3K27ac did not change with age, indicating that enhancer marking and co-accessibility do not necessarily go hand-in-hand (Figure S4). Taken together, integration of scATAC- with scRNA-seq data confirms that alterations in chromatin states are linked to gene expression differences. However, on a global level, we observed a disconnect between chromatin alterations and transcriptional output, suggesting some decoupling of chromatin states and transcription with age.

### Cellular heterogeneity in gene expression but not in chromatin states increases with age

The observation that co-accessibility and transcription were not correlated in general (Figure 3f) and the finding that scRNA-seq data did not fully identify the age of cells during cell type prediction (Figure 3c) suggested that there is a decoupling between chromatin architecture and steady-state levels of mRNA in ageing hepatocytes. To identify the underlying reason for this observation, we investigated the decoupling between chromatin and the transcriptome. We initially projected the available data on liver tissue from the Tabula Muris senis consortium as a UMAP, which was generated using either the 10x Genomics platform (droplet data) or using flow cytometry and Smart-seq2 (FACS data). Consistent with the outcome of the cell type prediction, the clustering based on scRNA-seq data did not resolve the different age groups, while it clearly separated the different liver tissue cell types (Figure 5a,b). This effect can also be observed in a PCA (Figure 5c) and remained apparent when focussing exclusively on hepatocytes (Figure 5d). Such a lack of ageing signature during clustering can be observed in other reports as well^53, 54^. A few studies have linked organismal and cellular ageing to transcriptional variability and cell-to-cell gene expression heterogeneity^3, 55^. Thus, we wondered if an increase in cell-to-cell heterogeneity would potentially mask any underlying transcriptional ageing signature in scRNA-seq data. For simplicity, we initially focused on the major cell type of the liver, hepatocytes. First, we fit a linear model for the first three PCs with age, taking into consideration biological independent experiments in the form of mouse identity (two mice per condition) as a confounding factor (Figure S5a). We calculated the adjusted R^2^ to quantify how well each PC explained age (Figure 5e) which remains under 25%. However, the noise explained as a sum of residual squares significantly increased in old cells (Figure 5f, Methods). Together, this analysis indicated that on a global level, only ∼22% of the expression patterns (variance) could be explained by age and the heterogeneity of hepatocytes strongly increased with age. To assess this in all other liver-resident cell types, we fit a linear model taking into consideration cell type as an additional variable. Noise increased in all cell types with the notable exception of B cells, which showed a decrease in noise with age (Figure S5b).

**Figure 5:**
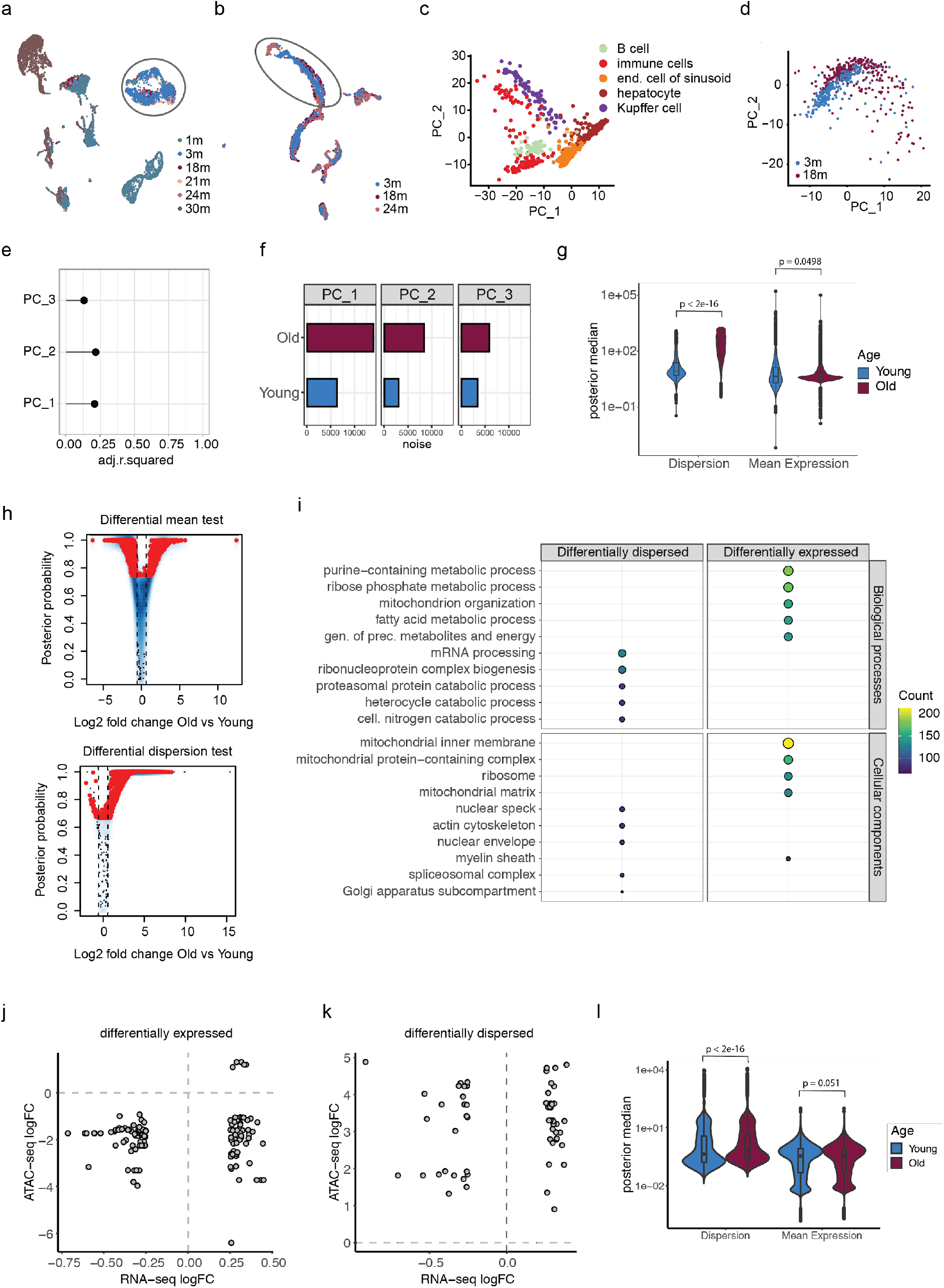
Transcriptional variability in hepatocytes increases with age. UMAP projection of Tabula Muris Senis (TMS) male a) 10x Genomics-based and b) FACS data coloured by age. Hepatocytes are marked with a circle. c) PCA projection of TMS male FACS data coloured by cell types and d) hepatocytes coloured by age. e) Adjusted R^2^ of the linear model fit of age and mouseID with the first three PCs (see Figure S4A). f) The sum of residual squares for each linear model fit to the first three PCs colored by age g) The posterior medians of mean expression (mu) and over-dispersion (delta) parameters estimated by a regression model from BASiCS coloured by age. P-values were calculated using a Welch Two Sample t-test. h,i) Log2 fold changes (x-axis) of significantly differentially expressed and over-dispersed genes with the differential accessibility log2 fold changes (y-axis) measured from scATAC-seq data. j) scATAC-seq gene activity matrix was used to estimate mean expression (mu) and over-dispersion (delta) parameters using a regression model from BASiCS colored by age. P-values were calculated using a Welch Two Sample t-test. k) Differential expression and variability was determined between young and old hepatocytes. For each gene, the difference in mean expression and over-dispersion is estimated as log2 fold-change (x-axis) and the posterior probability (y-axis) where the red highlighted genes are significantly differentially expressed or dispersed. l) Top biological processes (upper panel) and cellular components (lower panel) enriched in the differentially dispersed (left) and differentially expressed (right) genes (Supplementary Table 6).

To identify genes that contributed to the age-dependent increase in noise, we used a regression model implemented within BASiCS ^56, 57^. As shown in our previous analysis, the overall distribution of mean expression remained similar while dispersion was observed to be significantly higher in old cells, as suggested by the median posterior estimates of young and old hepatocytes (Figure 5g). By this means, we were able to compare the differential variability between young and old cells for the genes with similar mean expression. The differential test obtained 5545 and 6537 genes significantly differentially expressed and dispersed, respectively (Supplementary Table 6).

The expression difference was found to be nearly symmetrical, with 2448 up- and 3097 genes down-regulated in old cells. With respect to variability, virtually all (6487 of 6537) genes showed significantly higher dispersion in old cells (Figure 5h). We further filtered the differentially over-dispersed genes for minimum 5% detection rate in each age group and mean overall expression 5 to account for low expression or detection rate, which retained 2020 significantly over-dispersed genes in old hepatocytes. Strikingly, differentially expressed and dispersed genes showed a clear functional separation with respect to pathways affected (Supplementary Table 6). GO enrichment analysis showed that an increase in cell-to-cell variability was associated with genes involved in mRNA processing RNP complex biogenesis (Figure 5i), indicating that genes involved in gene expression regulation showed a particular increase in variability with age. On the other hand, differentially expressed genes were enriched for GO terms that deal with metabolic processes, translation and mitochondrial organisation (Figure 5i). This finding was supported by the similar results from KEGG pathway enrichment (Figure S5c). When compared to the bulk RNA-seq data (Figure S1a,b), the over- dispersed genes contributed to 27.06% of the differentially expressed genes and 32.4% of the dispersed genes overlapped with the global ageing genes^16^. Finally, we carried out the same analysis with the TMS Droplet and FACS data of female hepatocytes at 3 and 18 months of age. We observed very similar effects in ageing female hepatocytes regardless of the scRNA-seq approach (Figure S5d). The overall dispersion was higher in aged cells and additionally, the functional network was found to be the same with >75% of the genes also overlapping between the datasets.

The cell type prediction of scATAC-seq data with the TMS scRNA-seq data did not resolve cell type age. Because of this apparent decoupling of chromatin state and transcription, we next correlated differential expression and dispersion in RNA with differential accessibility in chromatin. We did not observe any correlation between RNA expression changes and chromatin states (Figures 5j,k). Finally, we decided to investigate if chromatin itself would show an increase in dispersion with age and performed BASiCS on the gene activity matrix of the scATAC-seq data (for details see Materials and Methods). In contrast to scRNA-seq, we did not observe an apparent increase in dispersion with age (Figure 5i), suggesting that chromatin states are less heterogeneous than the transcripts. This difference in the dispersion might also be one underlying reason for the observed decoupling between RNA and chromatin states in the single-cell data. In summary, we observed a very strong increase in cell-to-cell variability over age in the regulatory gene network, potentially masking mean expression differences and hindering the ageing signature from being detected by single-cell gene expression analysis.

## DISCUSSION

The question of how the direct microenvironment of a cell within a tissue affects the ageing trajectory has not been extensively explored. A few studies investigated the role of the microenvironment, particularly on the fate of tissue-resident stem cells, in which age-dependent perturbations of e.g. the vascular niches trigger the loss of functional hematopoietic stem cells and osteoprogenitors^58^. Indeed, general attrition of vascularisation has been recently reported occurring in multiple organs, including the liver^22^ indicating that tissue microenvironments experience profound alterations with age. This is in line with the observation that ageing is accompanied by a decline in blood flow in the liver^59^. Given the importance of the vascular system in setting up the division of labour of hepatocytes, the liver represents an ideal tissue to address the consequences of tissue organisation and location on one cell type.

Next to the insights into the connection of micronenvironmental changes and metabolic as well as epigenomic changes in the ageing liver, the data represent a valuable resource for researchers interested in liver organisation. While the scATAC-seq data will allow the interrogation of chromatin states in most of liver-resident cell types, the spatial transcriptomics data will mostly give insight into hepatic functions as the hepatocyte are dominating the transcriptional profiles on the spots. However, manual inspection of marker cell types indicates that also the spatial data can be used to interrogate non-parenchymal cells, particularly Kupffer, endothelial and stellate cells (Figure S6).

The most apparent and macroscopic alteration with ageing to liver physiology is the accumulation of large LDs in a zonated pattern, with the bulk of LDs being localised in hepatocytes around the central vein of the liver lobule. Using spatial transcriptomics we explored the age-dependent changes that occur within the central to portal axis of the liver lobule. Interestingly, we identified members of the Cide gene family to be predominantly upregulated in the central area of the liver lobule. Cidea, Cideb and Cidec are important regulators of LD dynamic and growth. Indeed, an increase in expression of Cidec has been shown to lead to growth of LDs^60^, suggesting that the increase in Cidea and Cidec expression might be one underlying reason for the increase in LD size with age. The changes in Cidea expression are also encoded in the epigenome. As our scATAC-data provided enough resolution to investigate zonation- and age-dependent differences, we could show that the locus encoding for Cidea is remodelled with age and co-accessibility increased. The presence of H3K27ac indicated that during ageing, an intronic enhancer is associated with the pericentral increase of Cidea expression in hepatocytes. Such an increase of expression in Cidea and Cidec has also been linked to the development of hepatic steatosis^61, 62^ and prolonged hepatic lipid storage can lead to metabolic dysfunction in the liver and inflammation. Ultimately, this development can lead to advanced forms of non-alcoholic fatty liver disease (NAFLD)^63^. Thus, it is no surprise that ageing is the most common cause for the progression of NAFLD.

Interestingly, the strong accumulation of large LDs in the pericentral region did not go hand-in-hand with major chromatin rearrangements. In fact, pericentral hepatocytes from young and old liver were called to belong to one topic only, indicating that their chromatin states were similar. On the other hand, young and old periportal hepatocytes differed sufficiently enough in their chromatin state to be enriched for different topics. Our lipidomic and spatial transcriptomic analysis might provide an explanation for the apparent difference in chromatin architecture in periportal hepatocytes. Cardiolipins and ubiquinones were altered strongly in aged cells. Together with measurements of mitochondrial respiratory capacity, the results indicated a change in efficiency of the electron transport chain and thus, ATP production, particularly in the periportal region of the liver. As periportal cells are exposed to high levels of oxygen due to their position close to the artery, they would usually rely on respiration. A decrease in vasculature^22^ and blood flow^59^ might therefore have stronger consequences on metabolic status in these hepatocytes than pericentral ones. A profound change in mitochondrial metabolism might have direct consequences on chromatin. Indeed, several studies have already connected changes in mitochondrial metabolism with alterations in chromatin structure^64–66^. In support of the hypothesis that a decrease in vasculature leads to changes in liver oxygenation, the spatial transcriptomics highlighted hypoxic signalling changed with age, specifically in the central region of the lobule.

Spatial transcriptomics and the scATAC data both showed a clear signature of ageing in hepatocytes. On the other hand, we noted that the scRNA-seq provided by the Tabula Muris Senis consortium^9^ was not able to cluster cells based on ageing. Even in hepatocytes, age explained only around 25% of the variance in the data. This low impact of ageing on clustering in scRNA-seq data can also be observed in other tissues in the Tabula Muris senis dataset and in a few studies that were published recently^53, 54^. In addition, while cell type prediction of the scATAC data worked well using scRNA-seq, different ages were distributed fairly evenly across the young and old hepatocyte clusters. This indicated a global decoupling of chromatin and RNA, which we confirmed by correlating changes in accessibility and gene expression. RNA-sequencing measures the steady-state level of mRNA, thus the technology would not be able to distinguish between changes in the synthesis and post-transcriptional regulation of mRNA^67^. Intriguingly, genes involved in post-transcriptional processing are among the top- dispersed genes, suggesting that this layer of gene expression regulation might be deregulated and more stochastic with age. One part of this layer would be mRNA splicing and indeed, there have been several reports over the last years that the process of splicing is strongly impacted by age and might itself contribute to ageing^68–70^. Totally unexplored as of now is the role of mRNA stability and storage with age. The decoupling of chromatin state and transcription is reminiscent of the decoupling of mRNA and protein levels with age^71^. Together, these data suggest that there is a progressive loss of cohesion between the different layers of gene expression that might contribute to the ageing process.

## MATERIALS AND METHODS

### Mice

C57BL/6N male young (3-4 months) and old (18-22months) old mice were bred and maintained in the mouse facility of Max Planck Institute for Biology of Ageing following ethical approval by the local authorities. The lights are controlled by timers and set a photoperiod of 12 hours of light from 6 am until 6pm (with a 15min twilight period). The room temperature is 22 +/- 2°C and the relative humidity 50 +/-5 %. All mice were fed with a standard diet ssniff M- Haltung, phyt.-arm (gamma irradiated).

### Immunohistochemistry

Livers were excised post-mortem and fixed directly into 4% PFA for 24hrs at 4°C, washed twice with 1XPBS, embedded into paraffin blocks and cut into 5μm sections. For Oil-Red-O staining and spatial transcriptomics, freshly-dissected liver tissues were frozen in Tissue-Tek OCT compound (Sakura) and cut into 7μm and 10μm cryosections, respectively.

For IHC stainings, sections of paraffin-embedded samples were deparaffinised by immersion of the slides into the following buffers; 20 min in Xylol, 2 min. 100% EtOH, 2 min. 96% EtOH, 75% EtOH and 1x PBS and washed three times with H_2_O for 5 min each. Endogenous peroxidase was quenched by immersion for 15 min in peroxidase blocking buffer (0.04 M NaCitrate pH 6.0, 0.121 M Na2HPO4, 0.03 M NaN3, 3% H_2_O_2_). After three washes with tap water, slides were subjected to heat-induced epitope retrieval with 10 mM NaCitrate, 0.05% Tween-20, pH 6.0, washed 5 min with 1X PBS, blocked 60 min with Blocking buffer + 160 µl/ml AvidinD and incubated with primary antibodies diluted (1:200 Plin2) in blocking buffer + 160 µl/ml Biotin overnight at 4°C. After three 5 min washes with PBST the samples were incubated with the secondary antibody 1:1000 diluted in blocking buffer for 1 h at room temperature, followed by three 5 min washes with PBST and incubation for 30 min with 1x PBS + 1:60 Avidin D + 1:60 Biotin. After three 5 min washes with PBST the samples were stained with 1 drop of DAB chromogen in 1 ml Substrate buffer, washed with 1X PBS and counterstained with Hematoxylin for 4 min, washed with tap water and distilled H2O and dehydrated 1min in each buffer; 75% EtOH, 96% EtOH, 100% EtOH, Xylol and mounted with Entellan.

### H&E staining

Following deparaffinization, slides with tissues washed with distilled and tapped water and stained with Ηematoxylin for 5 min, followed by 5 washes in tapped water and staining with Eosin Y for 3 min, followed by 3 washes with tap water, dehydration and mounting in Entellan.

### Oil-red-O and Sirius Red staining

Oil-Red-O and Sirius Red staining were used to visualize neutral lipids and collagen, respectively, and were performed according to standard procedures. Oil-Red-O staining was performed on 7-μm-thick frozen liver sections that were fixed in 4% paraformaldehyde for 10 min, followed by staining with 0.3% Oil-Red-O (Sigma) in isopropanol/water (60:40 vol/vol) for 15min. Sirius red was performed on deparaffinized liver sections that were incubated for 1h at RT in Picro Sirius Red solution (ab150681, Abcam), followed by washes in acetic acid and alcohol solutions.

### RNAscope 2.5 HD Duplex

Liver tissue was placed in a cassette, fixed in 4%paraformaldehyde (PFA) dissolved in phosphate-buffered saline (pH 7.4) for 24hrs at 4°, washed twice with 1XPBS, and embedded into paraffin blocks. 7μm thick sections were processed as described below. Detection of *Cyp2f2* (Cat No. 451851), *Alb* (Cat No. 4437691), *Cyp2e1* (Cat No. 402781-C2) and *Glul* (Cat No. 426231-C2) mRNA was performed using a chromogenic *in situ* hybridization technique (RNAscope™ 2.5 HD Duplex Assay, Advanced Cell Diagnostics) according to the manufacturer’s instructions. RNAscope® 2.5 Duplex positive control probes PPIB-C1 and POLR2A-C2 (Cat No. 321651) were processed in parallel with the target probes. All incubation steps were performed using the ACD HybEz hybridization system (Cat No. 321462). Sections were mounted on SuperFrost Plus Gold slides (ThermoFisher), dried at RT, briefly rinsed in autoclaved Millipore water, air-dried, baked at 60°C for 1hrs and deparaffinized. Afterward, slides were treated with hydrogen peroxidase for 10 min. and submerged in Target Retrieval (Cat No. 322000) at 98.5-99.5°C for 30 min, followed by two brief rinses in autoclaved Millipore water. A hydrophobic barrier was then created around the sections using an ImmEdge hydrophobic barrier pen (Cat No. 310018). Sections were incubated with Protease Plus (Cat No. 322330) for 30 min. The subsequent hybridization, amplification and detection steps were performed according to the manufacturer’s instructions (2.5 HD Duplex Detection kit (Chromogenic), Cat No. 322500). Sections were counterstained with 50% Hematoxylin staining and mounted with VectaMount permanent mounting medium (Cat No. H-5000).

### Microscopy

Immunohistochemistry, stainings and RNA scope images were taken using a Nikon Eclipse Ci microscope, with a colour camera.

### Liver perfusion and flow cytometry

Livers were dissociated using the Miltenyi liver perfusion kit (beta-test version) following the manufacturer’s instructions. The isolated hepatocytes were washed two times with staining buffer (1x PBS, 2mM EDTA, 0.5%BSA) and 1-7million hepatocytes were stained with 1:50 FcX, 1:100 PE-anti-E-cadherin, 1:100 APC-anti-CD73 for 1hr at room temperature. Cells were washed two times with staining buffer, cells were filtered through a 100um strainer dead cells were excluded with DAPI. Cells were sorted using a BD FACSARIA IIIU or Fusion Cytometer and 130um nozzle. The data were analysed using the BD FACSDiva and FlowJo softwares.

### Mitochondrial function measurement

Mitochondrial function was evaluated by measuring the Oxygen Consumption Rate (OCR) with the Seahorse XFe96 Extracellular Flux Analyzer (Agilent). XFe96 cell culture plates were coated with Collagen-I (40 μg/ml) overnight at 4°C and then washed 2x with 1X DPBS before 6,000 murine primary hepatocytes were seeded onto each well. Cells were cultured overnight in DMEM+GlutaMAX containing 10% FBS and 1x PenStrep under humidified conditions at 37°C with 5% CO_2_. Cells were washed 2x with assay media composed of XF DMEM medium (pH 7.4) supplemented with glucose (10 mM), pyruvate (1 mM) and glutamine (2 mM). Cells were cultured in assay media and incubated for 1h at 37 °C in a non-CO_2_ incubator. The Seahorse XF Mito Stress test was used to measure the OCR response after the sequential injection of oligomycin (1.0 μM), FCCP (1.0 μM) and Rot/AA (0.5 μM), according to the manufacturer’s instructions. The data were normalised to cell numbers.

### Genomic DNA extraction and qPCR for mitochondrial content

Cells were trypsinised and genomic DNA was extracted using the NucleoSpin Tissue XS, Micro kit for DNA (REF 740901.50). Real time PCR was performed with primers specific to the cyto-b mitochondrial locus (fw: TCCGATATATACACGCAAACG, rv: ATAAGCCTCGTCCGACATGA) and results were nomalised to total genomic DNA using primers for actin promoter locus (fw: TGCCCCATTCAATGTCTCGG, rv: ATCCACGTGACATCCACACC).

### mRNA extraction and qPCR for Cyp2f2 and Glul expression

To verify the relative abundance of expression of the respective markers of the sorted cells, CD73+ pericentral and E-cadherin+ periportal cells were isolated with flow cytometry (see methods above) from 3 individual (1 young and 2 old) mice and mRNA was extracted with the Dynabeads™ mRNA DIRECT™ Purification Kit (61011 Thermo Fisher Scientific). Reverse transcription was performed with the Maxima H Minus Reverse Transkriptase (EP 0751 Thermo Fisher Scientific) and the cDNA was used for qPCR with primers for Cyp2f2 (fw: CTTCCTGATACCCAAGGGCAC, rv: CTGAGGCGTCTTGAACTGGT) and Glul (fw: CCACCGCTCTGAACACCTT, rv: TGGCTTGGACTTTCTCACCC). The results were normalised to Actin expression (fw: ACCGGTGCAGAGACATTGGAGTTCAAC, rv: GTCGACTCAGATCCCGAGGCAGAGTC).

### Lipidomics

#### Lipid extraction from liver tissue samples or liver duct organoids

For the lipidomic analysis of liver tissue, 20 mg of snap-frozen tissue samples were homogenised using pre-cooled (liquid N_2_) metal balls (5 mm diameter) in a Qiagen Tissue Lyser for 1 min at 25 Hz. The pulverized tissue was resuspended in1 ml pre-cooled (−20°C) extraction buffer (MTBE (methyl tert-butyl):MeOH 75:25 [v:v]), containing 0.2 µL of EquiSplash Lipidomix as an internal standard. The re-suspended samples were homogenised for additional 5 min at 15 Hz in the TissueLyser.

After efficient tissue lysis, the samples were incubated for additional for 30 min on a thermomixer at 1500 rpm and at 4°C. To remove precipitated material from the samples, the Metal balls were removed and all samples were centrifuged for 10 min at 4°C and21.000 x g. The supernatants was transferred to a new tube and 500 µl H_2_O:methanol 3:1 [v:v] were added before further incubating the extracts for additional 10 min at 1500 rpm and 15°C on a thermomixer. After this final incubation step the polar and lipid phases were separated in a 10 min centrifugation step at 16.000 x g and 15°C. . The upper phase, MTBE-phase was transferred to a new tube and stored with the obtained insoluble pellets at -80°C for lipidomic analysis and protein extraction and quantification (BCA).

#### Liquid Chromatography-High Resolution Mass Spectrometry-based (LC-HRMS) analysis of lipids

The stored (−80°C) lipid extracts were dried in a SpeedVac concentrator before analysis and lipid pellets were resuspended in 200 µL of a UPLC-grade acetonitrile: isopropanol (70:30 [v:v]) mixture. Samples were vortexed for 10 seconds and incubated for 10 min on a thermomixer at 4°C. Re-suspended samples were centrifuged for 5 min at 10.000 x g and 4°C, before transferring the cleared supernatant to 2 ml glass vials with 200 µl glass inserts. All samples were placed in an Acquity iClass UPLC sample manager at 6°C. The UPLC was connected to a Tribrid Orbitrap HRMS, equipped with a heated electrospray ionization (HESI) ion source (ID-X, Thermo Fischer Scientific).

Of each lipid sample, 1 µl was injected onto a 100 x 2.1 mm BEH C_8_ UPLC column, packed with 1.7 µm particles. The flow rate of the UPLC was set to 400 µl/min and the buffer system consisted of buffer A (10 mM ammonium acetate, 0.1% acetic acid in UPLC-grade water) and buffer B (10 mM ammonium acetate, 0.1% acetic acid in UPLC-grade acetonitrile/isopropanol 7:3 [v/v]). The UPLC gradient was as follows: 0-1 min 45% A, 1-4 min 45-25% A, 4-12 min 25-11% A, 12-15 min 11-1% A, 15-18 min 1% A, 20-18.1 min 1-45% A and 18.1-22 min re-equilibrating at 45% A. This leads to a total runtime of 22 min per sample.

The ID-X mass spectrometer was operating either for the first injection in positive ionization mode or for the second injection in negative ionization mode. In both cases, the analyzed mass range was between m/z 160-1600. The resolution (R) was set to 120.000, leading to approximately 4 scans per second. The RF lens was set to 60%, while the AGC target was set to 250%. The maximal ion time was set to 100 ms and the HESI source was operating with a spray voltage of 3.5 kV in positive ionization mode, while 3.2 kV were applied in negative ionization mode. The ion tube transfer capillary temperature was 300°C, the sheath gas flow 60 arbitrary units (AU), the auxiliary gas flow 20 AU and the sweep gas flow was set to 1 AU at 340°C.

All samples were measured in a randomized run-order and targeted data analysis was performed using the quan module of the TraceFinder 4.1 software (Thermo Fischer Scientific) in combination with a sample-specific in-house generated compound database. Peak areas of each peak were normalized to the internal standards from the extraction buffer and to either the fresh weight of the tissue or the protein concentration of the organoids.

### Spatial transcriptomics

#### Tissue and library preparation

Liver specimen from 2 young and 2 old mice were cryopreserved and sections of 8 mm x 8 mm x 10μm specimens. Libraries were prepared using the Visium Spatial Gene Expression solution from 10x Genomics using 30 minutes permeabilization time. Libraries were prepared according to the manufacturer’s instruction and sequenced on an Illumina NovaSeq 6000. Sequencing data was initially quality controlled and pre-processed using the 10X Genomics CellRanger framework.

#### Dimensionality reduction and individual analysis of datasets

Young and old liver tissue slides were analyzed individually in R (V. 4.0.0) using the Seurat package (V. 4.0.4)^41^. Count matrices were normalized and scaled using the *SCTransform* function with standard parameters. Relative gene expression visualization of known hepatic pericentral and periportal marker genes on the spots of the tissue slides was performed with the *SpatialFeaturePlot* function.

#### Dataset integration

To assess batch effects between tissue slides, we merged the processed slides using the *merge* function and normalized and scaled without any further batch correction. Principal component analysis for Figure 2B was performed on the 2000 most variable features. The top 50 genes associated with the first principal PCA component were visualized with the *VizDimLoadings* functions and intersected with the hepatocyte specific aging genes list from Ref. 14^16^. Integration of young and old liver tissue slides was performed in a stepwise manner as an integration of all datasets together would remove all potential differences between young and old datasets. First, the pre-processed young and old tissue slide datasets were integrated separately per age group using canonical correlation analysis described in ^15^. Second, both combined datasets were merged and filtered for spots to have at least 1000 and at most 7000 genes expressed. Subsequently, the joined count matrix was scaled and normalized together using the *NormalizeData* and *ScaleData* function.

#### Dimensionality reduction of integrated datasets

We performed principal components analysis on the preprocessed data (*RunPCA* function). The first 10 principal components covered most of the data set’s variance, and were considered a good approximation to the data as assessed by an elbowplot (*Elbowplot* function). The first 10 principal components, therefore, served as input to UMAP for further dimension reduction and visualization. Known canonical liver zonation marker genes were visualized with the *Featureplot* function.

#### Differential expression testing between young and old liver tissue slides

Differential expression testing was done by using the *FindMarkers* function. Genes had to show at least an average log_2_-fold change of ±0.25 to be considered for testing. Testing was performed using the *MAST* library by^17^. Bonferroni correction was applied for multiple testing adjustments of p-values. Go-term enrichment analysis for combinatorial categories was performed with the *enrichGO* function from the *clusterProfiler* library^72^. Results were summarised using REVIGO (http://revigo.irb.hr/)^73^. Heatmap visualization of genes from categories of interest was done with pheatmap^74^. Genes of GO-terms were extracted from the *org.Mm.eg.db* library ^75^. Log_2_ fold changes were calculated using the *FoldChange* function.

### Cytoscape

The Cytoscape^18^ apps ClueGo^19^ and iRegulon^23^ were used to calculate gene ontology networks and transcription factor predictions, respectively. All differentially expressed genes in old (Supplementary Table 4) were used as input for all analysis. ClueGo parameters were as follows: Biological Pathways were selected as ontologies and only pathways with pV ≤ 0.001. GO Tree Interval was between 6 and 12. Cluster #1 was set at 2 minimum genes that represented 5% of genes, while the network connectivity was set at 0.4. iRegulon was run using Mus musculus MGI symbols using the following motif collection: 10k (9712PWMs). Putative regulatory regaion as well as motif ranking database were set as 20kb centered around TSS. NES scores for all TFs reported were > 4.

### Liver tissue preparation for scATAC-seq

Liver nuclei (n=4) were prepared from frozen tissue specimens by crushing and dounce homogenising the tissue in 1 ml EZbuffer (SIGMA) (20 strokes with loose and a tight pestle, respectively) and spun 5 min at 300 g. The pellet was incubated on ice for 20 min in EZ-buffer supplemented with DNAseI NEB M0303S (4 units/ml) and 1X DNAseI buffer. Equal volume of EZ-buffer was added and samples were spun 5 min at 500 g and incubated again 10 min on ice in EZ-buffer supplemented with DNAseI NEB M0303S (8 units/ml) and 1X DNAseI buffer. Equal volume of EZ-buffer was added, and samples were spun 5min at 500g, resuspended in NSB (1087.5 µl 1XPBS, 5.5µl 2% BSA, 1.5 µl RNase Inhibitor) and filtered 3 times through a 0.22 µm strainer. For scATAC-seq, 100,000 nuclei were resuspended in 50 µl tagmentation mix (10X Genomics)).

### scATAC-seq library preparation and sequencing

scATAC-seq targeting 4000 cells per sample was performed using a beta version of Chromium Single Cell ATAC Library and Gel Bead kit (10x Genomics, 1000110) according to the manufacturer’s instructions. Libraries were then pooled and loaded on an Illumina NovaSeq sequencer and sequenced to 18,904 median reads per cell for the young dataset and 21,139 median reads per cell for the old dataset. Sequencing data was initially quality controlled and pre-processed using the 10X Genomics CellRanger framework.

### scATAC-seq analysis of young and old liver tissue

Region accessibility count data were analyzed using the *cisTopic* library *(V. 3.0*^39^. Cells without any accessible regions were removed, leaving 4838 cells from young mice and 3361 cells from old mice. We included 117,290 regions into our analysis that were accessible in at least one cell. The latent Dirichlet allocation model was learned by the *runWarpLDAModels* function for topic numbers ranging from 2 to 15 topics. An appropriate number of topics for our data was selected as the topic number with the highest second derivative of the likelihood function. This was the case for 6 topics, and all downstream analyses use the LDA model learned for 6 topics. Non-linear dimensionality reduction by UMAP was performed for visualization purposes only by applying the built-in *runUmap* function in cisTopic to the topic-distributions of all cells. Topic defining regions were derived via the *getRegionsScores*- and *binarizecisTopics*-function. GO-term and transcription factor motif analysis of the topic defining regions was done using *rGREAT (V.1.22.0)*^76^ and *RcisTarget (V.1.10)*^77^. Transcription factor motifs shown in Fig 3F and Fig 6B were downloaded from the *JASPAR* database (http://jaspar.genereg.net).

To check the robustness of the cisTopic results, we performed a complementary analysis of the ame data with *Signac (V.1.0)*^40^. The cell region count matrix was normalized using the term frequency-inverse document frequency (TF-IDF) normalization method from the *Signac* library (*RunTFIDF*). Initial linear dimensionality reduction was performed with singular value decomposition (*RunSVD*). As recoded in the Signac workflow, the first component of the singular value decomposition was excluded from all downstream analyses as it was highly correlated with the sequencing depth. Non-linear dimensionality reduction (UMAP) for Fig. S3A+B was generated via the *RunUMAP* function. The dimensions 2 to 35 were used as input for the algorithm.

#### Differential accessibility testing

We employed the *FindMarkers* function in the logistic regression framework of^78^ to test for regions that were differentially accessible between young and old hepatocytes, respectively, between periportal and pericentral hepatocytes. We considered only regions detected in at least 5% of the cells for testing. P-values were Bonferroni adjusted to account for multiple testing.

#### Cell type annotation

Our celltype annotation is based on the imputed gene activity of known liver cell marker genes from *CellAtlas*^79^. To calculate the imputed gene activities, fragments mapping to gene bodies or promoter regions of genes (Up to 2 kb upstream of a gene) were summed up using the *GeneActivity* function and subsequently normalized via the *NormalizeData* function from Signac. Periportal and pericentral cell populations were annotated based on the gene activity of *Cyp2e1* and *Cyp2f2* genes.

#### Cell classification via canonical correlation analysis

Tabula Muris Senis ^9^ droplet data were preprocessed as described in the respective section in the manuscript and filtered for cells for male individuals between 3 and 30 months of age. Transfer anchors were determined using the *FindTransferAnchors* function. Cell labels from the tabula Muris droplet dataset were used as provided in the metadata. Cell labels for the scATAC-seq dataset were predicted with the *TransferData* function. For details, see^15^.

#### Construction of Cis-regulatory networks

Co-accessibility scores for the interaction network of the *Cidea* locus were predicted with the *Cicero* library^51^. Reduced dimension coordinates of cells were based on the UMAP projection from *cisTopic*. Connections of co-accessible loci were inferred for young and old hepatocytes separately.

### Bulk RNA-seq data processing and analysis

The TMS bulk RNA-seq data was analysed as described above by directly using the count matrix provided (https://doi.org/10.6084/m9.figshare.8286230.v1). We only used the data from male mice of the age 3 and 18 months.

### scRNA-seq data processing and analysis

#### Preliminary processing of TMS data

We downloaded metadata and raw count tables from Tabula Muris Senis consortium for liver FACS and droplets methods. The TMS FACS and droplets data was filtered for genes expressed in at least 3 cells, cells containing minimum 250 genes and 2500 counts for droplets while 500 genes and 5000 UMIs for the FACS data. The filtered count matrix was processed using Seurat (4.0.4)^41^ with default parameters as per suggested pipeline using ‘NormalizeData’, ‘FindVariableFeatures’, ‘ScaleData’, ‘RunPCA’, ‘RunUMAP’, ‘FindNeighbors’ and ‘FindClusters’ functions. The feature and PCA/UMAP plots generated in this manuscript are through Seurat plotting functions.

#### Linear model fit of the principal components

We obtained the cell embeddings for each principal component from the processed Seurat objects. The input parameters are principal components, age and animal identity of the cells. The linear model for only hepatocytes was fitted using the ‘lm’ function in R (4.0.1) as lm(PC_n ∼ Age + Mouse.id). The model estimates and predictions were extracted using the R package broom (https://CRAN.R-project.org/package=broom). The model fit with cell types was done in the same manner with “cell type” as an additional factor for cell identity. We tested the increase in noise for significance with 10,000 permutations and compared the actual variance in the old and young cells. This test gave p-values of 0.0002, 0.018 and 0 for the first 3 PCs respectively.

#### Differential expression and dispersion analysis

The differential analysis was performed using the BASiCS package^56, 57^. Posterior estimates were computed using a Markov chain Monte Carlo (MCMC) simulation with 20,000 iterations and burn-in period 10000 with a regression model. We used BASiCS to detect differentially expressed and differentially variable genes between old and young hepatocytes. For changes in mean expression between ages, we use the ‘BASiCS_TestDE’ function with EFDR cutoff 0.1. Only genes with no change in mean expression were considered for interpreting changes in variability. We filtered genes with the detection rate of 0.5 in each age and mean overall expression of 5.

Obtained sets of genes from each differentially expressed and variability were further subjected to Gene Ontology Biological Processes enrichment analysis using the ‘enrichGO’ function from clusterProfiler (3.14.3) R package^72^. To remove the redundancy of enriched terms, we used the ‘simplify’ function from clusterProfiler with the default parameters. The pathway enrichment was performed using the ‘enrichPathway’ function from the ReactomePA R package (1.36.0)^80^.

## DATA AVAILABILITY

All sequencing data generated for this study is available at ENA under curation. H3K27ac for young and old mice was downloaded from https://www.ncbi.nlm.nih.gov/bioproject/?term=PRJNA28112781. Tabula Muris senis single cell data is available at: https://www.ncbi.nlm.nih.gov/geo/query/acc.cgi?acc=GSE1495909. Tabula Muris senis bulk RNA-seq data is available at: https://www.ncbi.nlm.nih.gov/geo/query/acc.cgi?acc=GSE1320409.

## AUTHOR CONTRIBUTIONS

Conceptualisation: C.N., S.P., N.K., A.T. and P.T.; Methodology: C.N., S.P., N.K., E.K., T.S., J.A.; Investigation: C.N., S.P., N.K., T.S., F.S., P.G., M.B., A.J.V., T.W. and E.K.; Formal Analysis: C.N., S.P., N.K. and P.G.; Supervision: A.T. and P.T.; Funding Acquisition: A.T. and P.T.; Project Administration: C.N., A.T., P.T.; Writing of Manuscript: P.T., with input from all authors

## ACKNOWLEDGMENTS

We would like to thank all members of the Tessarz and Tresch labs for continuous discussion. We are grateful to A. Schaefer and A. Pouikli for critical reading of the manuscript. We are indebted to the following core facilities of the MPI for Biology of Ageing for superb technical assistance: FACS and Imaging for help with FACS analysis, histology and microscopy, Metabolomics for lipidomic analysis and Comparative Biology for housing mice. Electron microscopy was performed at the Imaging Core Facility of CECAD, University of Cologne. scATAC-seq and spatial transcriptomics were performed at the Cologne Center for Genomics, University of Cologne, Germany. All other libraries were sequenced at the Sequencing Core Facility of the MPI for Molecular Genetics, Berlin, Germany. This work was funded by the Max Planck Society (to P.T. and T.W.), the Deutsche Forschungsgemeinschaft (DFG, German Research Foundation; project no. 415274764 (V.K. and F.S.), the BOOST program of the Max Planck Society (to C.N.) and the Deutsche Forschungsgemeinschaft (DFG, under Germany’s Excellence Strategy – EXC 2030 – 390661388) (to P.T.).

---

**Figure S1:**
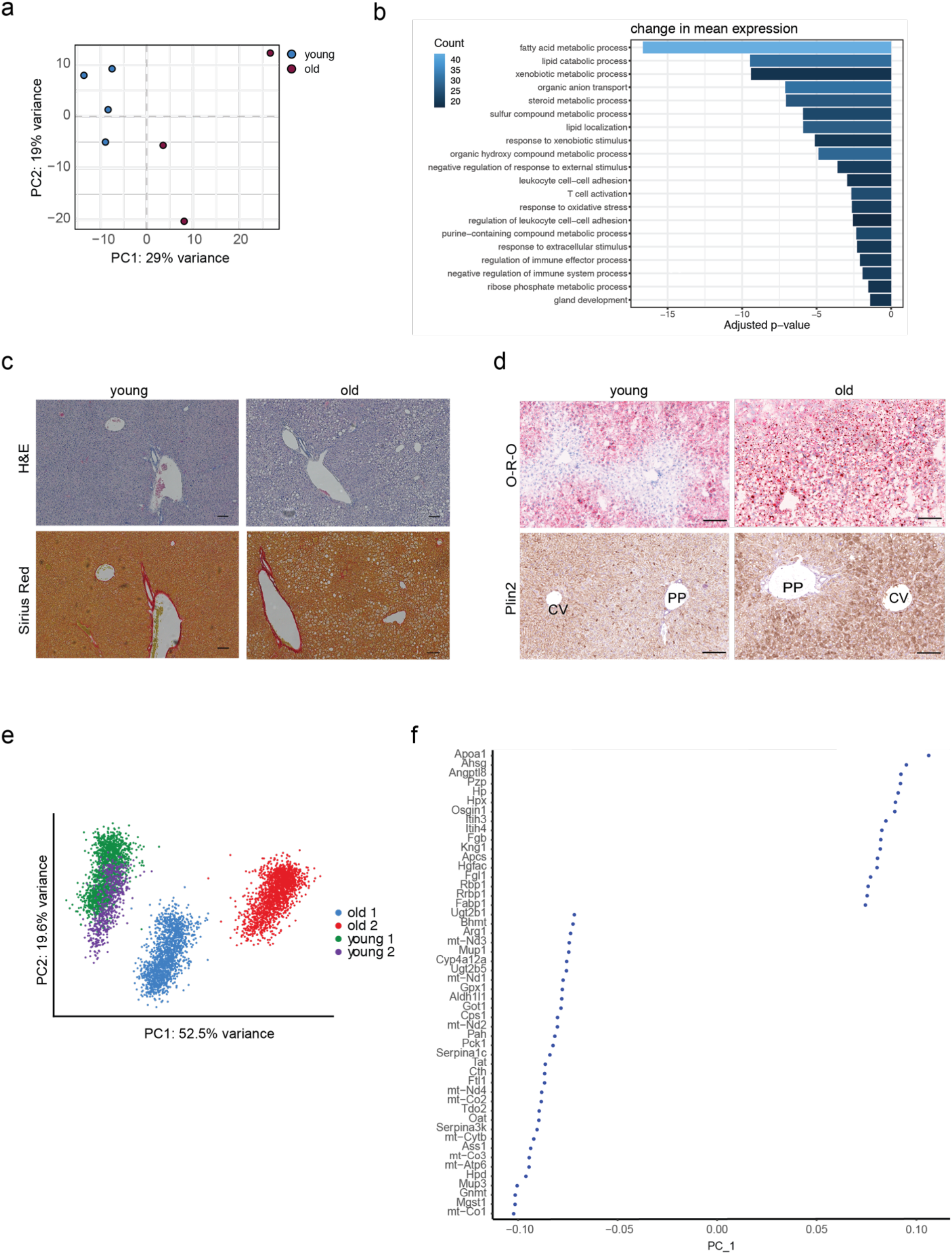
a) PCA projection of bulk RNAseq data ^9^ derived from young and old mouse livers. b) Differentially enriched pathways in the aged liver tissue derived from A (Supplementary Table 1). The colour scale represents the number of genes in each term. c) Representative images from H&E (upper panel) and Sirius Red (lower panel) stainings on liver sections from a young and an old mouse. Scale bar=100μm. d) Representative images of PP (periportal) and CV (central vein areas) of Oil-red-O (O- R-O, upper panel) and Plin2 immunostainings (lower panel) on liver sections from young and old mice. Scale bar = 100 µm. e) PCA plot of the spatial data after integration of the four datasets using canonical correlation analysis. Different colours represent the different samples. f) PC plot showing the top 50 genes that separate the ageing groups in Figure S1e.

**Figure S2:**
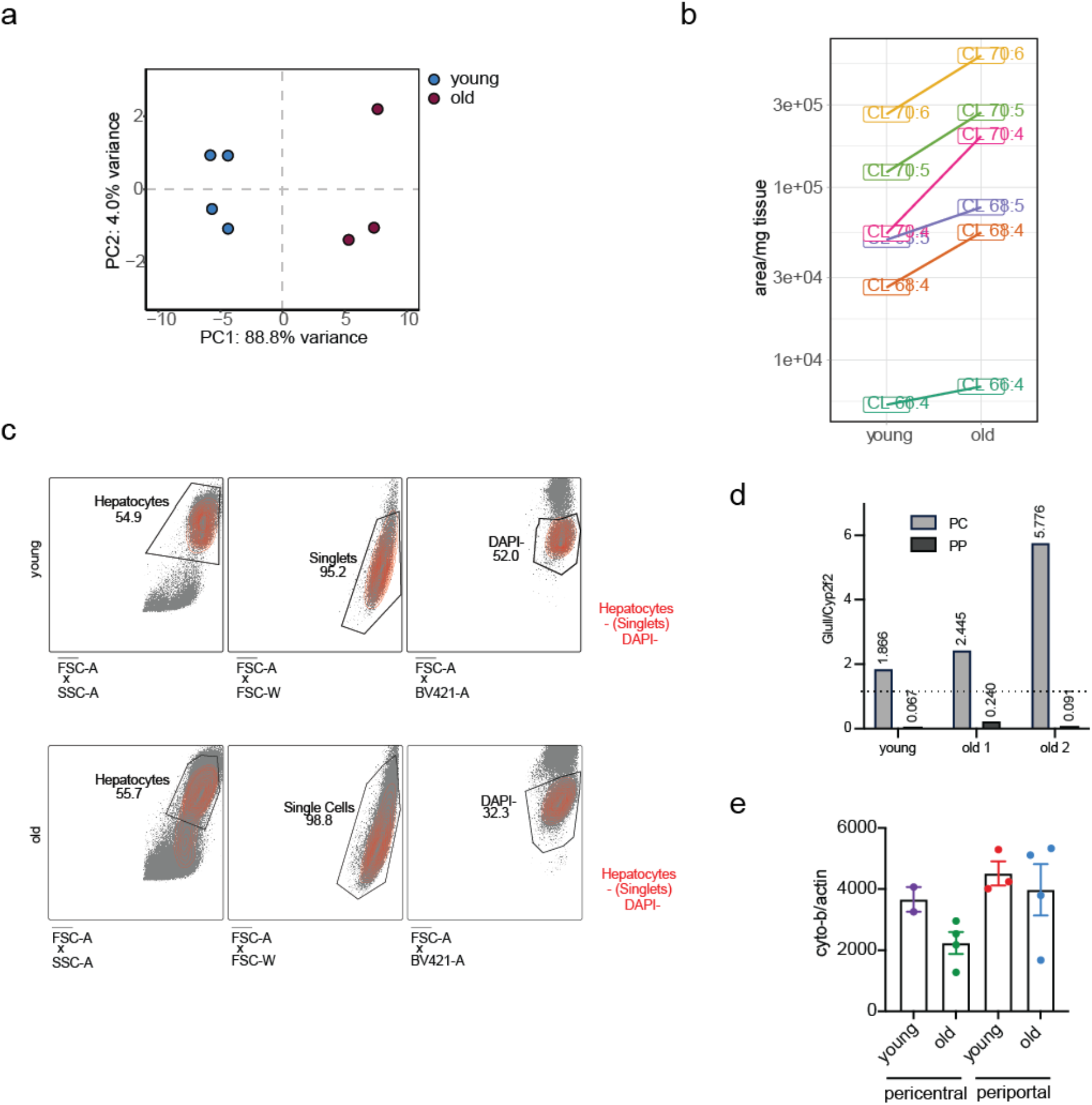
a) PCA of lipidomic data coloured by age. b) Changes of individual cardiolipins between young and old livers. c) Gating strategy for isolation of pericentral and periportal hepatocytes. d) qRT-PCR to validate the enrichment for pericentral and periportal hepatocytes based on expression ratios of Glul and Cyp2f2 levels. Shown are individual replicates for young and old mice (as indicated). e) Mitochondrial content was measured using primers against genomic copies of cyto-b and b-actin. Individual values are given as dots. Error bars represent the SEM.

**Figure S3:**
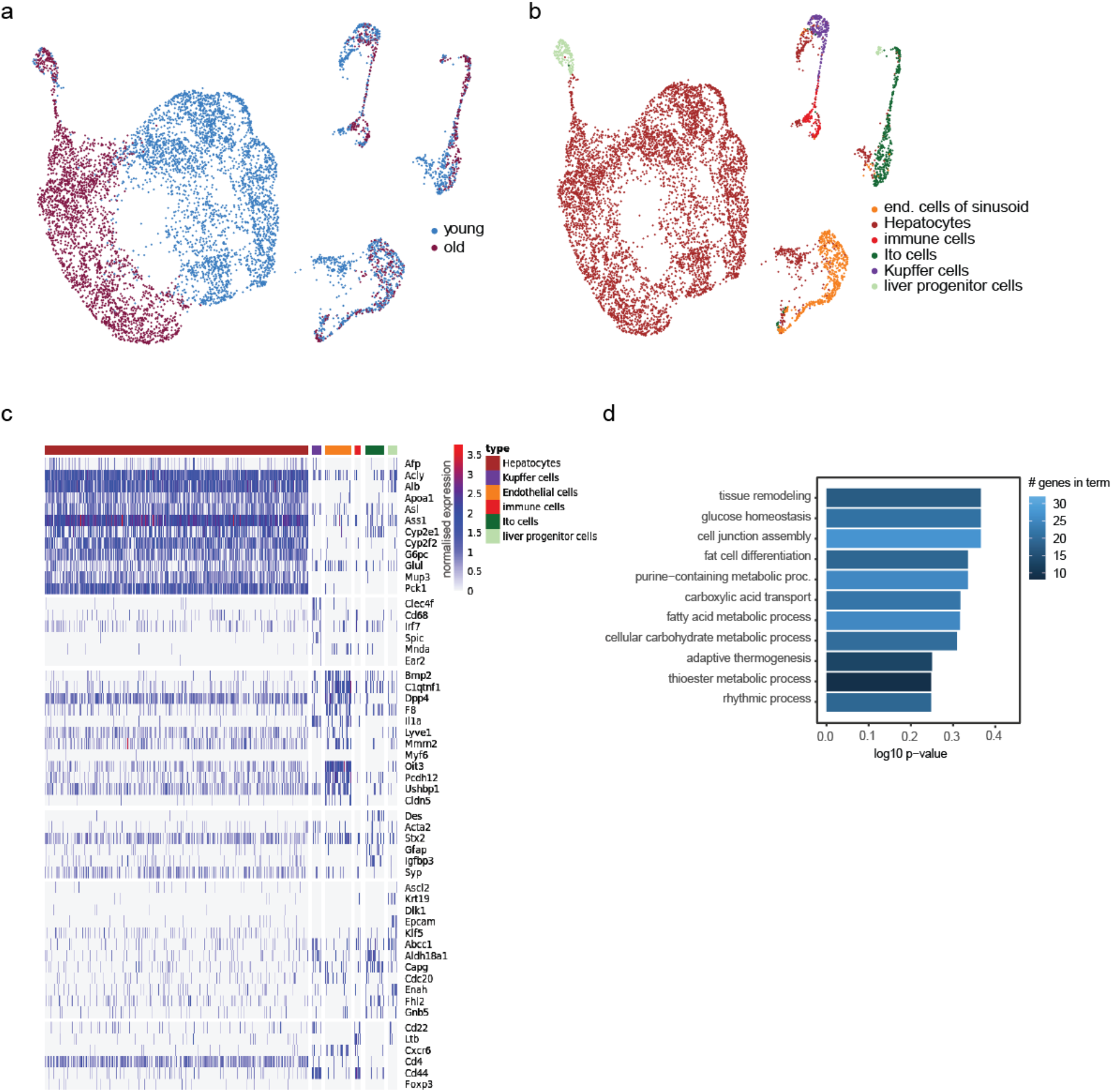
a) UMAP projection of scATAC-seq nuclei from young and old livers. Colour-coded are the different age groups identified using Signac. b) Same as in a). Colour coded are the different cell types, assigned by using marker genes from CellMarker. c) Heatmap showing the accessibility of marker genes in each assigned cell type of the scATAC-seq data. d) GO enrichment for genes found in differentially accessible loci in young vs. old hepatocytes (TSS+/- 3kb).

**Figure S4:**
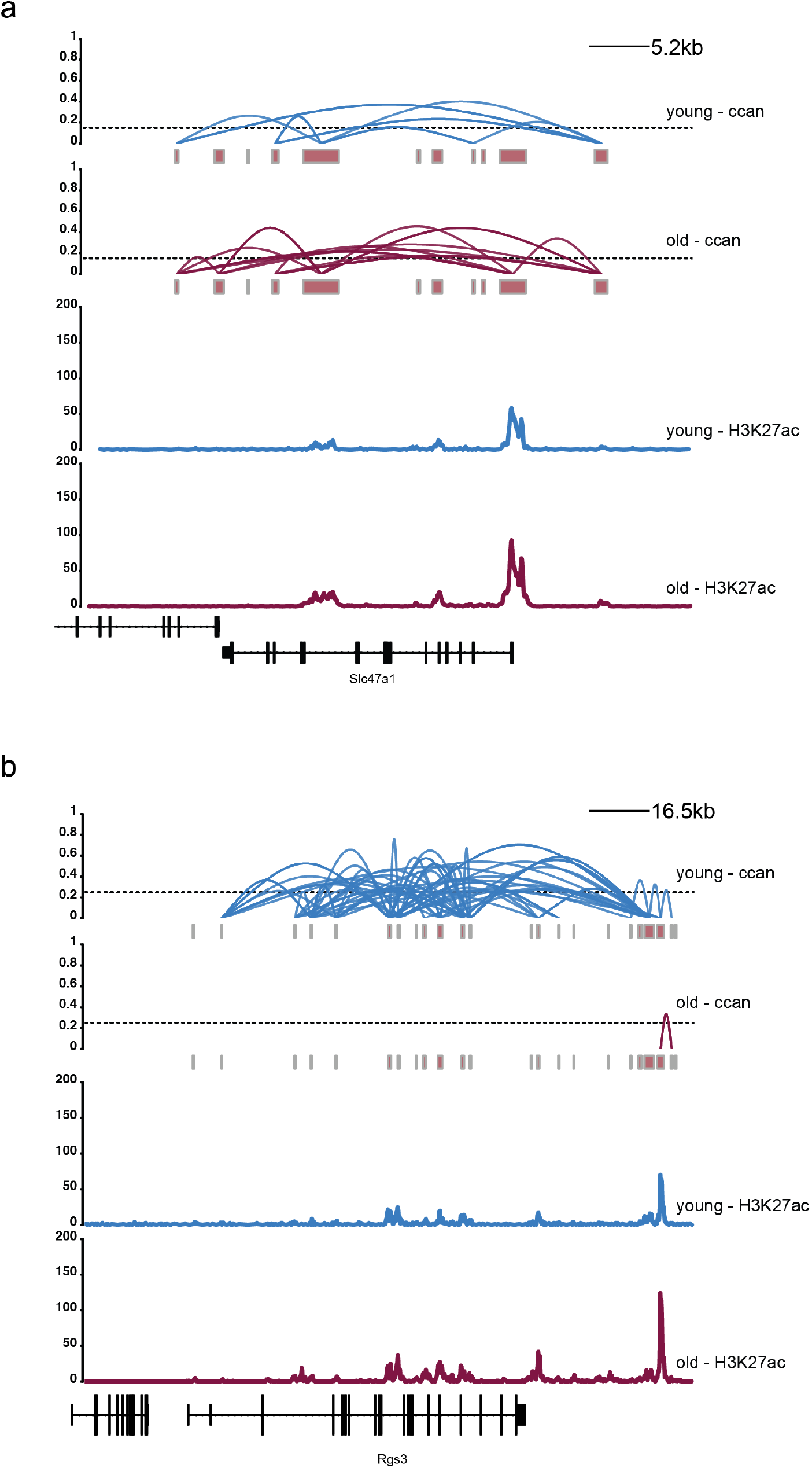
Ccan plots of loci identified to show increased (Slc47a1, a) and decreased (Rgs3, b) co-accessibility. H3K27ac tracks are shown to indicate potential enhancers.

**Figure S5:**
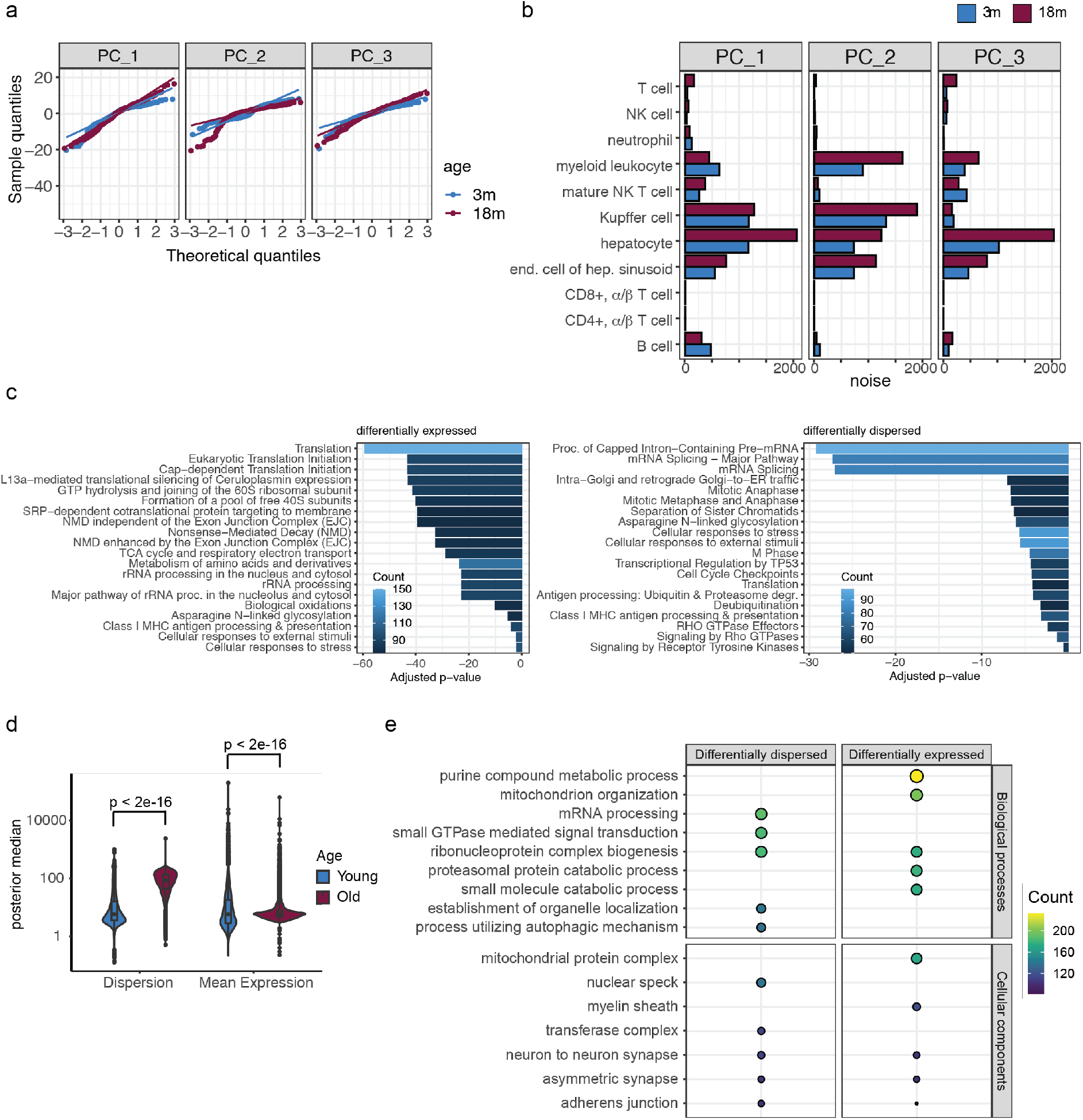
a) QQ-plot of the residuals from a linear model fit for the first 3 PCs with age. b) A barplot of the sum of residual squares (noise) for each linear model fit to the first 5 PCs with age and cell type coloured by age. c) The pathways enrichment for the differentially expressed (left) and differentially over-dispersed (right) genes. d) TMS FACS female data from age 3 and 18 was used to estimate mean expression (mu) and over-dispersion (delta) parameters using a regression model from BASiCS coloured by age. e) Top biological processes (upper panel) and cellular components (lower panel) enriched in the differentially dispersed (left) and differentially expressed (right) genes in the TMS FACS female dataset.

**Figure S6:**
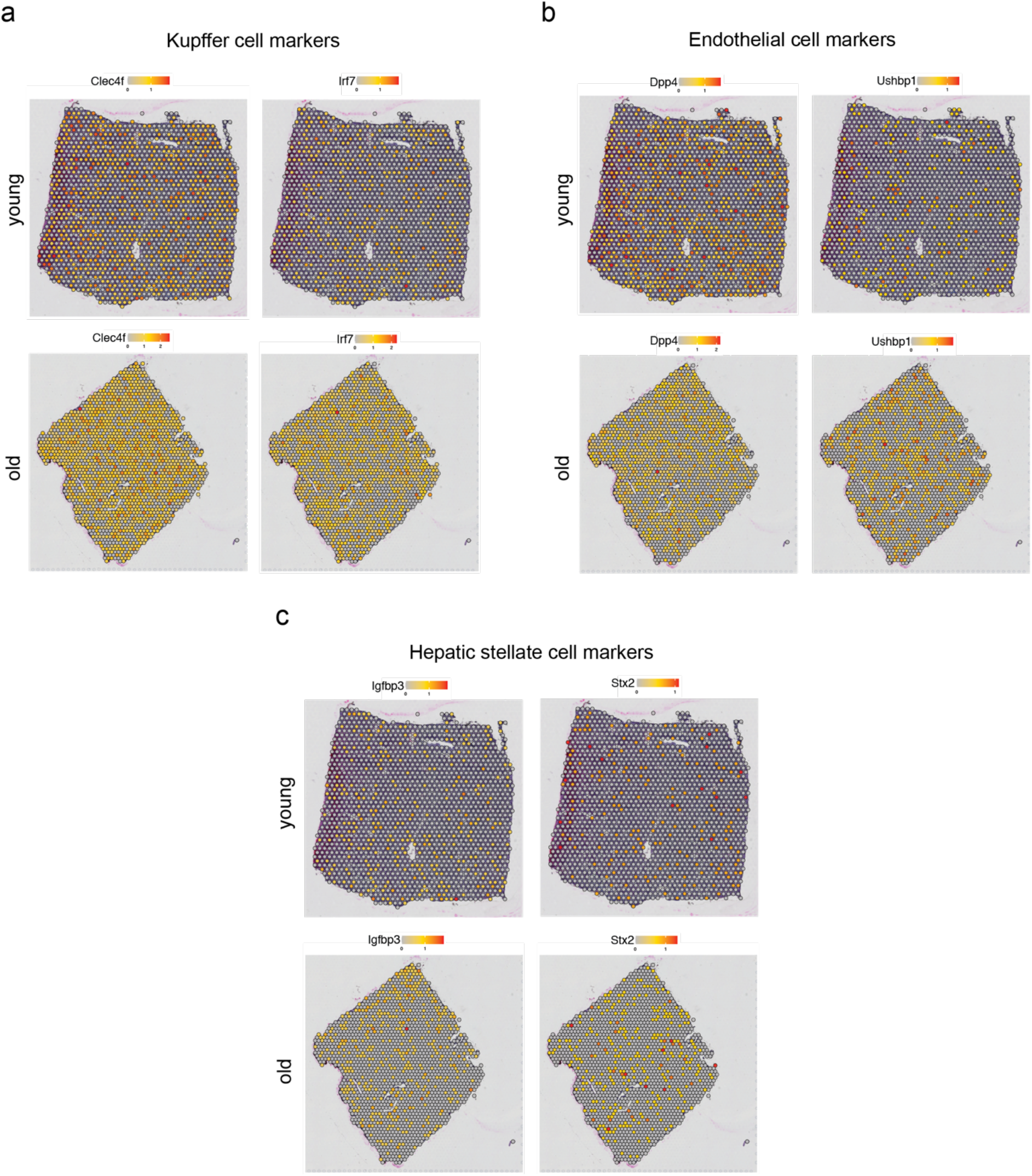
Representative plots showing expression levels of Kupffer cell (a), endothelial cell (b) and hepatic stellate cell (c) markers as indicated in young and old livers as determined by spatial transcriptomics. The colour gradient represents normalised gene expression.

